# Microinjection into the *Caenorhabditis elegans* embryo using an uncoated glass needle enables cell lineage visualization and reveals cell-non-autonomous adhesion control

**DOI:** 10.1101/406991

**Authors:** Yohei Kikuchi, Akatsuki Kimura

## Abstract

Microinjection is a useful method in cell biology, with which exogenous substances are introduced into a cell in a location- and time-specific manner. The *Caenorhabditis elegans* embryo is an important model system for cell and developmental biology. Applying microinjection to the *C. elegans* embryo had been difficult due to the rigid eggshell surrounding the embryo. In 2013, microinjection method using a carbon-coated quartz needle for the *C. elegans* embryo was reported. To prepare the needle, unfortunately, special equipment is required and thus a limited number of researchers can use this method. In this study, we established a method for the microinjection of drugs, dyes, and microbeads into the *C. elegans* embryo using an uncoated glass needle that can be produced in a general laboratory. This method enabled us to easily detect cell lineage up to adult stages by injecting a fluorescent dye into a blastomere. We also found a cell-non-autonomous control mechanism of cell adhesion; specifically, the injection of an actin inhibitor into one cell at the 2-cell stage enhanced adhesion between daughter cells of the other cell. Our microinjection method is expected to be used for broad studies and could facilitate various discoveries using *C. elegans*.

## Introduction

Microinjection is a useful method in cell biology. It can directly deliver substances prepared outside the cell to the inside of the cell with desired timing and to a desired location. For example, the role of microtubules and actomyosin in cell division were characterized by injecting inhibitors (O’Connell *et al.*, 1999; Strickland *et al.*, 2005), and the growth rate of the astral microtubules was measured by injecting oils or microbeads (Hamaguchi *et al.*, 1986). The *Caenorhabditis elegans* embryo is a major model system in cell biology. Sophisticated methods for gene manipulation enable researchers to express fluorescent proteins (Chalfie *et al.*, 1994), and the transparent embryonic cells permit observers to follow the processes of cell division and development under a microscope (Gönczy and Rose, 2005). Unfortunately, direct microinjection into the embryo has been considered difficult, as the embryo is covered by a rigid eggshell (Edgar *et al.*, 1994; McNally and McNally, 2005; Olson *et al.*, 2012; Marcello *et al.*, 2013; Stein and Golden, 2015).

To deliver substances into the *C. elegans* embryo, researchers perform microinjection into the gonads or soak the embryo in a solution containing the substance. Microinjection into the gonad is a popular approach to knock down gene function via RNAi or to obtain transgenic strains (Mello *et al.*, 1991). After microinjection into the gonad, and with sufficient time allotted, the substance will be incorporated into the embryo. Previously, using this method, microbeads or magnetic beads were introduced into the embryo to measure viscosity or forces inside the cells (Daniels *et al.*, 2006; Garzon-Coral *et al.*, 2016). However, this method is not efficient, as these substances will be diluted in the gonad; the time required for materials to be delivered to the embryos is also a disadvantage. The other method (soaking) is also difficult when using most substances as the eggshell acts as a permeability barrier. To enable these materials to penetrate the embryo from the outside (Strome and Wood, 1983; Schierenberg and Junkersdorf, 1992), permeability must be increased by knocking down genes such as *perm-1* (Carvalho *et al.*, 2011). It should be noted that the knockdown of *perm-1* causes embryonic lethality, and are thus invasive. Most importantly, with both methods—injection into the gonad and soaking—it is impossible to introduce substances into the embryo in a time- or location-specific manner.

In 2013, a method for direct microinjection into *C. elegans* was developed using a carbon-coated quartz needle (Brennan *et al.*, 2013). This report demonstrated that substances could be directly introduced into the embryo. Unfortunately, the carbon-coated quartz needle is difficult to obtain in an ordinary biology lab (including that of the authors of this paper), as carbon coating requires special equipment.

In this study, we succeeded in establishing a microinjection method for *C. elegans* embryos using an ordinary glass needle without special coating. With this method, we were able to deliver substances directly into the embryo in a time- and location-specific manner. We also evaluated the invasiveness of the method by quantifying the rate of cell division and hatching after injection. By injecting a fluorescent dye into a blastomere, this method enabled us to detect cell lineage easily, up to adult stages. We also found a cell-non-autonomous control mechanism of cell adhesion as follows: the injection of an actin inhibitor into one cell at the 2-cell stage enhanced the adhesion between the daughter cells of the other cell.

## Results

### Preparation of glass needles for microinjection

In an attempt to establish a method for microinjection into *C. elegans* embryos using ordinary glass needles, we prepared glass needles with a shape similar to the carbon-coated quartz needles used in the preceding research (Brennan *et al.*, 2013). By changing the input parameters of a micropipette puller (Sutter Instrument, P-1000), we prepared three types of glass needles with long and thin tip regions. We measured the outer and inner diameters of the tips of the glass needles by scanning electron microscopy (SEM) (Fig. 1A). The average inner diameters of the three types of the needles were 100 nm, 150 nm, and 657 nm, respectively, and we referred to each type as ‘*φ*100’, ‘*φ*150’, and ‘*φ*660’, respectively (Fig. 1B). The *φ*100-type had the smallest inner diameter, and was the most similar to the quartz needle among the three types. We also quantified the ejection volume of the glass needles using the method of the previous report (Brennan *et al.*, 2013). We ejected a fluorescent-dextran solution into glycerol (Fig. 1C) with a pressure of 1,000 hPa (14.5 psi) for 0.5 s, which was the minimum setting used to eject a reproducible volume with our equipment. The ejection volume of the *φ*100-type was 16.0 fl, which was comparable to that of the quartz needle in the previous study (Brennan *et al.*, 2013) (Fig. 1D). By assuming the long axis of the embryo is 50 μm and the short axis is 30 μm, the volume of the embryo was estimated to be 24,000 μm^3^. Therefore, the ejection volume was estimated to be about 0.1 % of the embryo volume. The ejection volume of the *φ*150-type was 33.8 fl. We were not able to quantify the ejection volume of the *φ*660-type (see Methods). As the ejection volume of the *φ*100-type was closest to that of the quartz needle, we used the *φ*100-type needle for subsequent experiments unless otherwise indicated.

**Figure 1.**
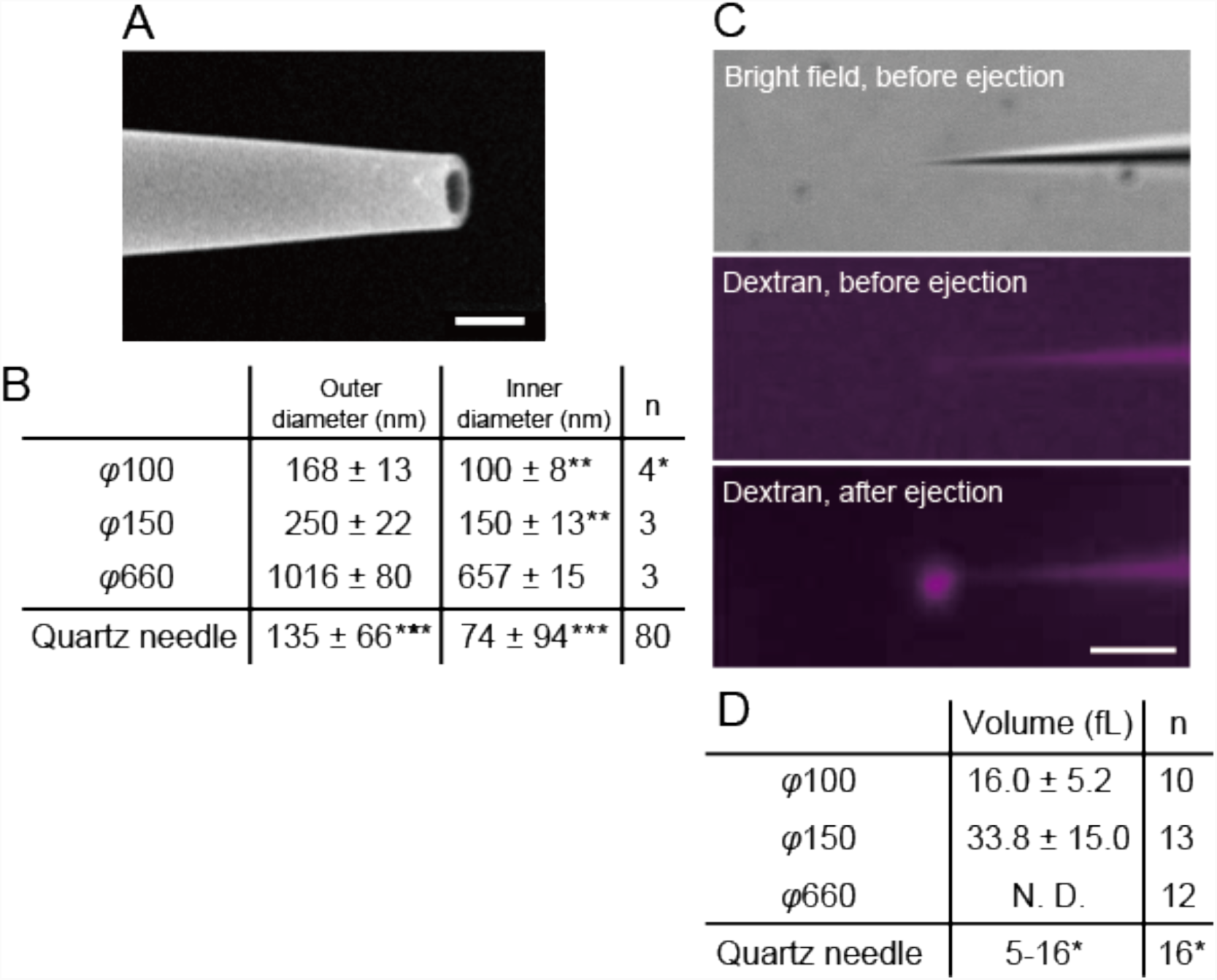
Observations of tip diameter of the glass needle and measurement of ejection volumes. (A) Examples of scanning electron microscopy (SEM) images of the tip of glass needles are shown. Bar, 1 μm. (B) Outer and inner diameters of the glass needle quantified based on the SEM images. * A needle with the measured outer diameter of 353 nm was excluded as it was an apparent outlier possibly due to the breakage of the tip during sample preparation for the SEM measurement. ** represents estimated values calculated based on the measured values of the outer diameter and theoretical ratio between the outer and inner diameter. *** in the bottom row represents values from previous research (Brennan *et al.*, 2013). (C) Bright field and fluorescence images of the glass needle before and after the ejection of dextran into glycerol to quantify the ejection volume. Bar, 10 μm. (D) Ejection volumes and percentages of the ejection volumes relative to the embryo volume. * in the bottom row represents values from previous research (Brennan *et al.*, 2013).

### Microinjection into the embryo with the glass needle was achieved by precise alignment of the micromanipulation system

As an initial attempt, we arranged the micromanipulation system in a similar manner to that used for the carbon-coated quartz needle (Brennan *et al.*, 2013). Using a holding pipette, an embryo was immobilized through its posterior pole. We attempted to insert the *φ*100-type needle, but it was unsuccessful as the needle tip slipped along the surface of the eggshell. Next, we fixed the embryo onto a silane-coated coverslip, which is used to fix starfish oocytes for microinjection (Kikuchi and Hamaguchi, 2012). This was also not successful as silane was not sticky enough to fix the position of the *C. elegans* embryo when it is pushed by the glass needle. Then, we returned to the approach using holding pipettes. To avoid the slippage of the needle on the eggshell, we found that two kinds of alignment were critical, namely, the ‘hold-needle alignment’ and ‘embryo-needle alignment’ (Fig. 2A, B). The ‘hold-needle alignment’ means that we precisely aligned the holding pipette and the glass needle in a straight line (‘hold-needle alignment’ in Fig. 2B). This hold-needle alignment is often used for microinjection into mammalian oocytes (Kimura and Yanagimachi, 1995). To achieve the alignment, a curved structure was introduced into both the holding pipette and the glass needle (Fig. 2A). The precise alignment was realized through adjustments under the microscope using fine micromanipulators. The ‘embryo-needle alignment’ indicates that the long axis of the embryos is perpendicular to the needle, such that the curvature of the eggshell at the point of injection will be minimum to avoid slippage. Additionally, a brake structure was introduced into the holding pipette for fine adjustments in aspiration pressure (Fig. 2A). With these efforts, we finally succeeded in inserting a *φ*100-type glass needle into the *C. elegans* embryo (Fig. 2A, enlarged image at the bottom).

**Figure 2.**
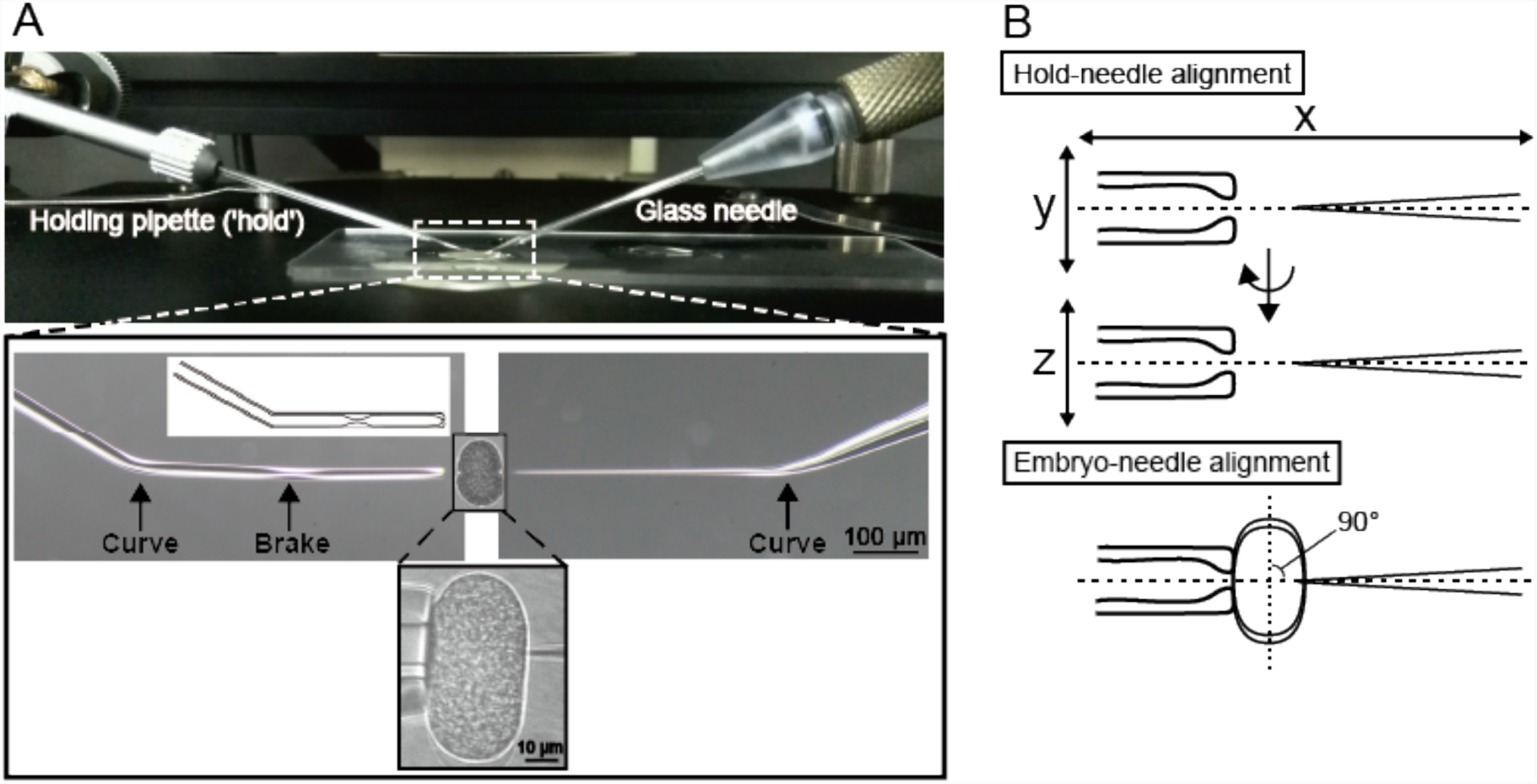
Micromanipulation system used in this study. (A) The upper image is a side view of the micromanipulation system on an inverted microscope. The holding pipette for immobilization of the embryo is on the left side and the glass needle for microinjection is on the right side. Lower images comprise an enlarged view of the region enclosed by the dashed line in the upper image. The bottom images are an actual example of micromanipulation. The tip region of both the holding pipette and the glass needle were precisely aligned in a straight line to insert the glass needle into the embryo. The pattern diagram shows the curve and brake structure within the holding pipette. (B) The ‘hold-needle alignment’ indicates the alignment between the holding pipette (‘hold’) and the glass needle, which should align straight from both x-y and x-z views. The ‘embryo-needle alignment’ denotes the alignment between the embryo and the needle. The long axis of the embryo should be perpendicular to the needle.

### Cells divided and embryos hatched after the microinjection procedure

We next evaluated the invasiveness of the microinjection procedure. We assessed the invasiveness of three steps individually, which were embryo immobilization, needle insertion, and buffer injection. We conducted the manipulation at the 1-cell stage, and scored the rate at which embryos entered the 4-cell stage and hatched. The embryos were maintained by the holding pipette until the 4-cell stage, and were then transferred to Shelton’s growth medium (SGM) (Shelton and Bowerman, 1996) and cultured overnight at 25 °C to score the rate of hatching.

First, the effect of embryo immobilization by the holding pipette was evaluated (Fig. 3A). Immobilization was conducted at various positions of the eggshell. All embryos divided twice to enter the 4-cell stage without noticeable delay and subsequently hatched (*n* = 12/12).

**Figure 3.**
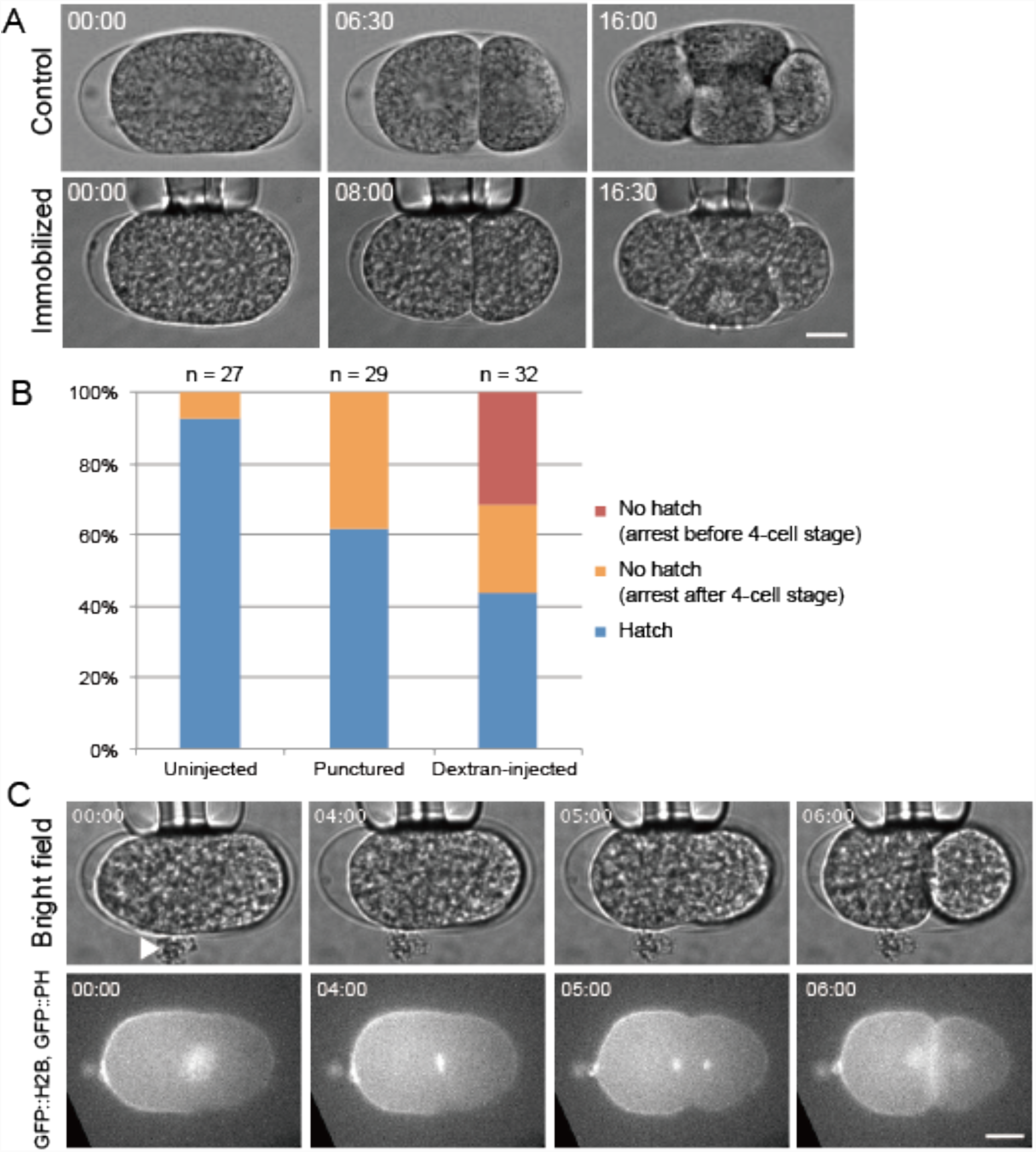
Micromanipulation of embryos and its effect on early embryogenesis. (A)Immobilization by the holding pipette did not disturb early *C. elegans* embryogenesis. Continuous images of control and immobilized embryo are shown. Immobilization of the embryo by the holding pipette did not disturb early embryogenesis (*n* = 12/12). The number in the upper left corner of the images shows an elapsed time after the start of observation (min:sec). The anterior is to the left. Bar, 10 μm. (B) Rates of completion of 4-cell stage and hatching in manipulated *C. elegans* embryos using glass needle methodology. Uninjected: embryo was immobilized by the holding pipette until the 4-cell stage with no needle insertion; punctured: a hole was made in the embryo by the glass needle but nothing was injected; dextran-injected: a hole was made in the embryo and Texas Red-dextran (MW = 3,000) in 0.8× egg buffer (EB) was injected. Embryos completing the 4-cell stage and those hatching were counted, and rates were determined. (C) Microinjection into 1-cell stage *C. elegans* embryos of CAL1041 strain using the glass needles. Continuous images of an embryo injected with Texas Red-dextran (MW = 3,000) in 0.8× EB. Cytoplasm leakage was often observed at the injection site when the glass needle was withdrawn from the embryo (arrowhead). However, embryogenesis progressed even if the cytoplasm leaked out after puncture via microinjection. The number in the upper left corner of the images shows the elapsed time after the start of observation (min:sec). Texas Red-dextran was used to monitor the success of substance delivery upon microinjection, and time-lapse imaging was conducted for bright field and green channels. The anterior is to the left. Bar, 10 μm.

Next, we evaluated the effect of inserting the needle into the cytoplasm (but not injecting) (Fig. 3B, Punctured). In some cases, when the glass needle was withdrawn, the cytoplasm leaked out of the eggshell. We noticed that the embryos with significant leakage failed cytokinesis, but if the leakage was small, the success rate of cytokinesis was high. We then set criteria that excluded embryos in which leakage occurred for more than 3 s after needle withdrawal from further analyses; 94% (*n* = 29/31) of embryos passed this criterion. Among the embryos that passed this criterion, 100% (*n* = 29/29) entered the 4-cell stage, and 62% (*n* = 18/29) of the embryos hatched (Fig. 3B, Punctured).

Finally, we evaluated the effect of injecting solution into the embryo. We injected Texas Red-dextran in 0.8× egg buffer (EB) into 1-cell stage embryos (Fig. 3B, Dextran-injected). After injection, 71% (*n* = 32/45) of embryos passed the criterion for leakage (i.e. 3-s). Among the embryos that passed the criterion, 69% (*n* = 22/32) of the embryos entered the 4-cell stage and 44% (*n* = 14/32) of embryos hatched. A successful example of cell division after the injection is shown in Fig. 3C and Video 1. GFP::H2B (histone) is a chromosome marker to monitor chromosome segregation, and GFP::PH (pleckstrin homology domain) is a membrane marker to monitor cytokinesis. When we used the *φ*150-type glass needles, 64% (*n* = 7/11) passed the leakage criterion. Among the embryos that passed this criterion, 57% (*n* = 4/7) reached the 4-cell stage and 29% (*n* = 2/7) hatched. When we used the *φ*660-type needle, massive leakage occurred and none of the embryos passed the criterion (*n* = 9). From the results, we concluded that we successfully established a method of direct injection into the *C. elegans* embryos using *φ*100-type needles; after the injection of dextran at the 1-cell stage, °70% of embryos that passed the criterion and divided twice to enter the 4-cell stage; moreover, greater than 40% hatched to become larvae. In conclusion, our method could be applied to analyze cell division and the development of embryos.

### Sizes of injectable substances

We next attempted to clarify the size range of injectable substances using this method. Using the three types of glass needles, we tested the ejection of substances of various sizes (dextrans or microbeads) into glycerol, or if they could be injected into 1-cell stage embryos (Table 1). The substances were loaded into the needle from the wider end. Dextran with MWs of 3,000 and 10,000 could be ejected into glycerol and into the embryo using all three types of the glass needles. Dextran with MW of 70,000 was ejectable only with the φ660-type needle, although the tips of the *φ*660-type needles were easily clogged with the dextran of this size. In this case, sonication and filtration treatments helped to avoid clogging.

**Table 1.**
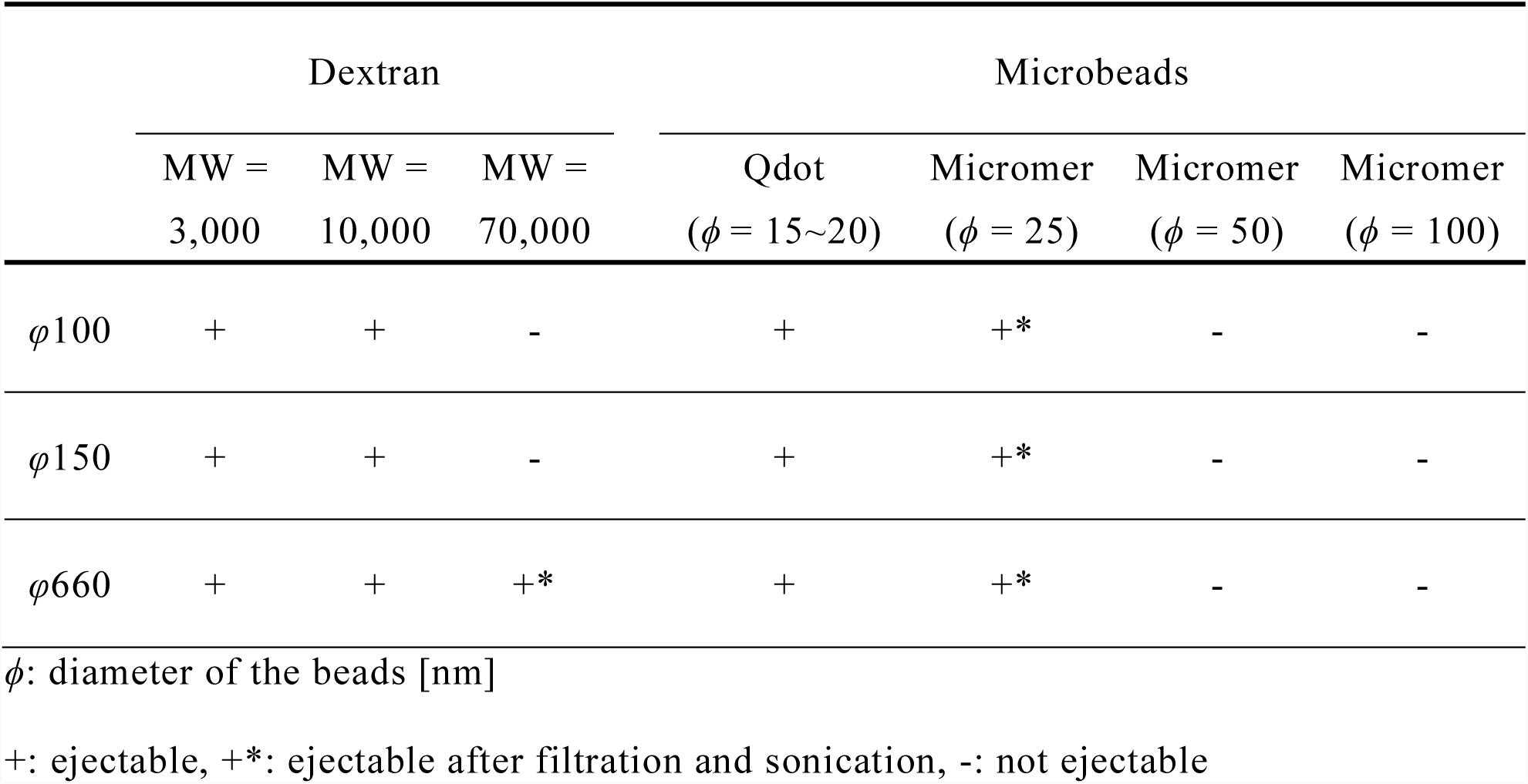
Relationship between inner diameters of the glass needle and size of ejectable substances

We next investigated microbeads of different sizes. Microbeads of 15–20 nm (in diameter) could be ejected into glycerol and into the embryos using all types of needles. In addition, 25-nm microbeads could also be ejected and injected if aggregations in the injection mixture were resolved before injection. Larger sized-microbeads (50 or 100 nm) could not be ejected or injected even after sonication, filtration, or dilution. In summary, dextran with a MW up to 10,000 and microbeads with a diameter up to 25 nm could be injected into the 1-cell stage embryo.

### Location-specific injection into 2-cell stage embryos

Thus far, we showed that using our method, substances can be injected directly into the *C. elegans* embryo with the desired timing (e.g. the 1-cell stage). We next attempted location-specific injection, which cannot be achieved by microinjection into the gonad or soaking (see Introduction). Dextran (MW = 3,000) was injected into one of the two cells (AB cell) at the 2-cell stage (Fig. 4). To achieve this, the holding pipette captured the eggshell near the AB cell (Fig. 4A). After cell division, at the 4-cell stage, fluorescent signals were observed only in the descendants of the AB cell (i.e. ABa and ABp cells), and not in the other cells (i.e. EMS and P2 cells) (Fig. 4B, Video 2).

**Figure 4.**
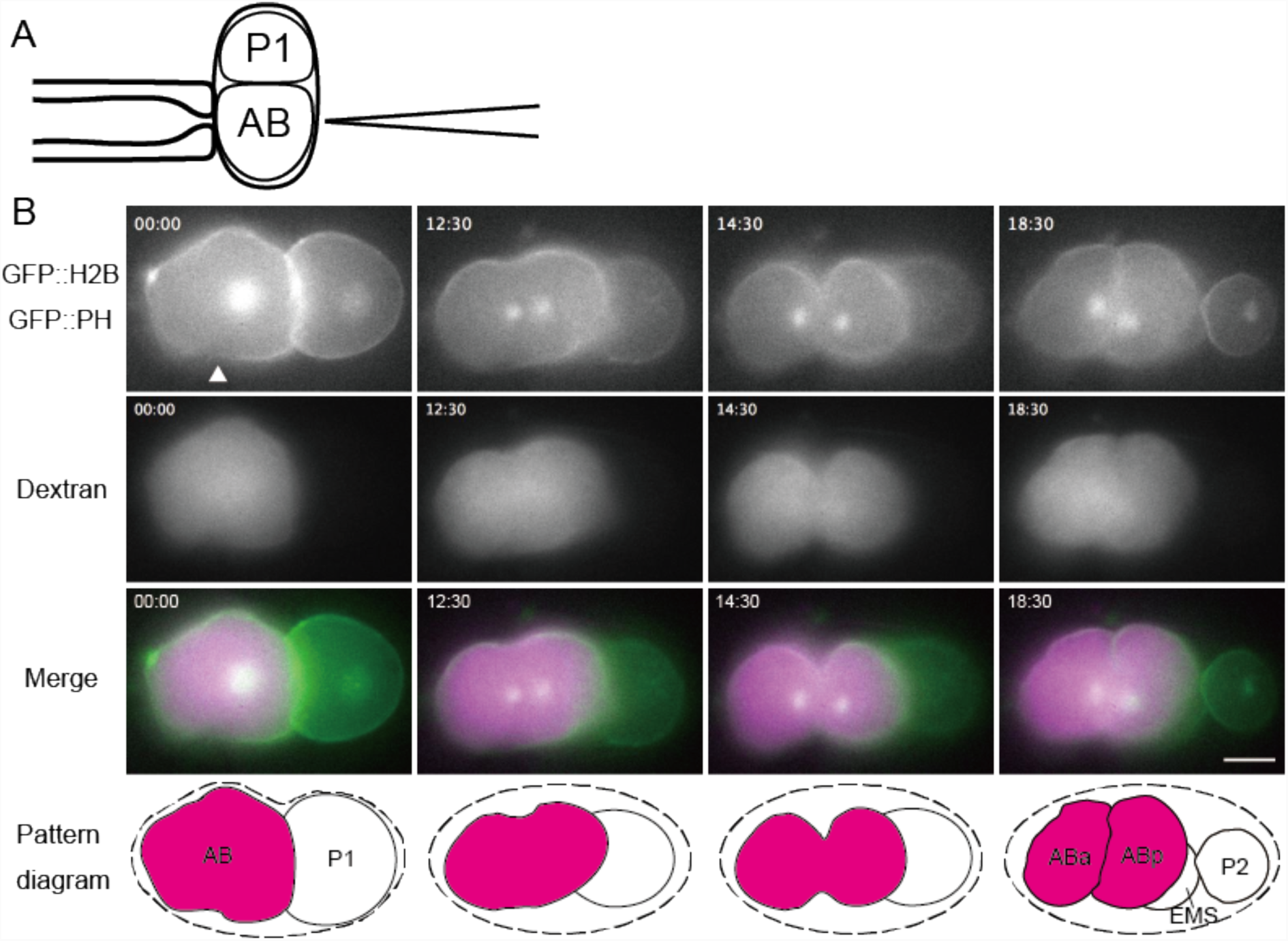
Dextran is distributed into a specific lineage of embryonic cells after microinjection. (A) Pattern diagram shows a capture position of the eggshell near the AB cell of the embryo and selective injection of dextran at the 2-cell stage. (B) Images show the dorsal view of the embryo. The number in the upper left corner of the images shows the elapsed time after the start of observation (min:sec). (00:00) Dextran was injected into the AB cell. The upper side of the AB cell was deformed by immobilization of the holding pipette and the lower side was deformed by injection of dextran. The arrowhead shows the dextran-injected site. (12:30) The AB cell had begun to divide. (14:30) AB cell immediately before completion of cell division. (18:30) At the 4-cell stage, fluorescent dextran was inherited only in the AB cell lineage. The bottom pattern diagrams are a summary of each stage. The dashed line indicates the presumed position of the eggshell. The anterior is to the left. Bar, 10 μm.

We further investigated whether the fluorescent signal could be detected in later stages. After the injection of fluorescent dextran into AB cells at the 2-cell stage, the fluorescent dextran signal was observed selectively in the AB cell lineage throughout embryogenesis (Fig. 5A–I). The AB cell lineage is known to differentiate primarily into ectodermal cells including hypodermis, neurons, and pharynx (Sulston *et al.*, 1983). At the ∼100-cell stage, approximately half of the cells of the injected embryo had dextran signal, and they occupied the surface of the embryo, as expected for ectodermal cells (Fig. 5I, Video 3). These embryos hatched after overnight incubation (*n* = 5/5). In hatched larvae, signals were observed in the hypodermis, neurons, and pharynx (Fig. 5J–O, arrows), but not in the germ cells derived from the P2 cell, as expected. Some signals were detected in the intestine, but they were thought to be autofluorescence as un-injected controls also had these signals (Fig. 5P–U). Surprisingly, the fluorescent dextran signal was not degraded or removed from the worm but the signals remained clear until the L4 stage (Fig. 5M). Some signals were detected even in adult worms in expected locations such as the pharynx (Fig. 5O). Our results indicate that the injection of fluorescent-dextran into a blastomere is an easy and powerful method to trace the cell lineage.

**Figure 5.**
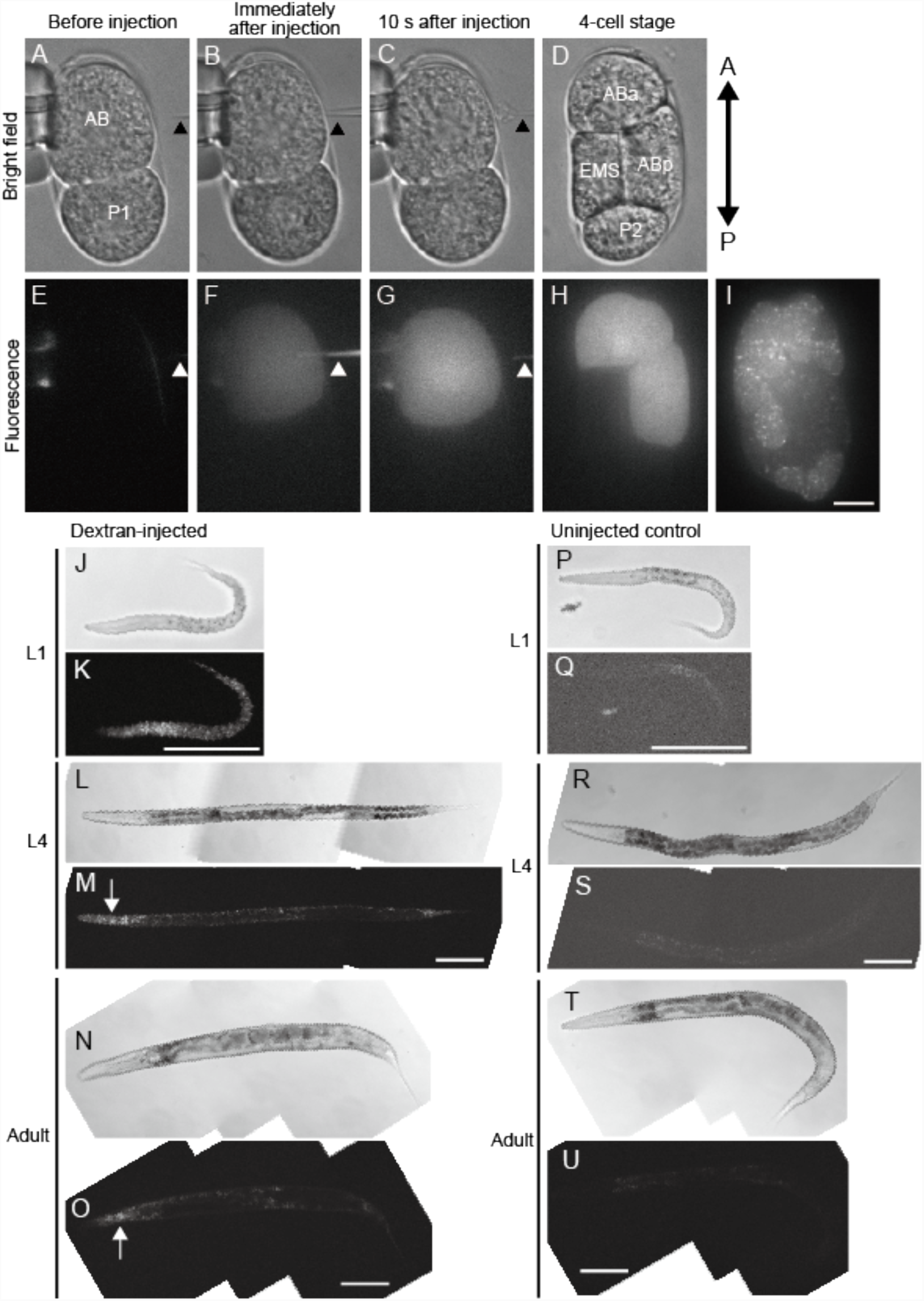
Injection and distribution of fluorescent dextran in the AB lineage. (A–H) Injection of fluorescent dextran into the AB cell. (A–D) Bright field. (E–I) Fluorescence. (I) An image at the °100-cell stage of the same embryo as in (A–H). Half of embryonic cells were observed to have a fluorescent dextran signal. (J–U) Images of fluorescent dextran-injected (J–O) and uninjected control (P-U) embryos at indicated stages. (J, L, N, P, R, T) bright field, (K, M, O, Q, S, U) fluorescence signal (561 nm excitation). Fluorescent signals were detected in the pharynx (arrows), hypodermis, and tail. (A–I) Bar, 10 μm. (J–U) Bars, 100 μm.

### A cell-non-autonomous effect of the actin cortex for proper cell arrangement

Location-specific injection enables us to inhibit the function of specific proteins in a desired cell. Such analysis can characterize cell-non-autonomous effects; if the inhibition of a protein in one cell affects the behavior of other cells, the effect is cell-non-autonomous. By injecting an actin inhibitor (Cytochalasin D) into the AB cell, we investigated the cell-non-autonomous effect of actin cortex integrity for proper cell arrangement at the 4-cell stage embryo.

The four cells at this stage are arranged into a diamond pattern in which the cells are attached to each other, except for between ABa and P2 cells (Fig. 6A, left). Our group has previously demonstrated that asymmetric attraction, in which the EMS cell tightly adheres to ABa and ABp cells, but weakly adheres to the P2 cell (Fig. 6A, middle), is important for the diamond arrangement even when the eggshell is deformed (Yamamoto and Kimura, 2017). It was not clear why the EMS cell adheres strongly to a set of cells (ABa and ABp) but not to the other (P2). A straightforward explanation is that the P2 cell has limited amounts of adhesive molecules (e.g. E-cadherin) and thus cannot strongly adhere to the EMS cell. However, when a P1 cell was isolated from an AB cell at the two-cell stage (after the eggshell was removed at the 1-cell stage), EMS and P2 (the daughters of P1) strongly adhered (Fig. 6A, right), which did not occur when the P1 cell was not isolated from eggshell-removed embryos (Fig. 6A, middle). The difference was quantified by measuring the length of contact area between EMS and P2 (Fig. 6B). The result suggests that the P2 cell has the potential to adhere strongly to the EMS cells, but that the potential is suppressed in normal conditions. From this result, we hypothesized that the suppression is caused by adhesion between EMS and ABa/p cells.

**Figure 6.**
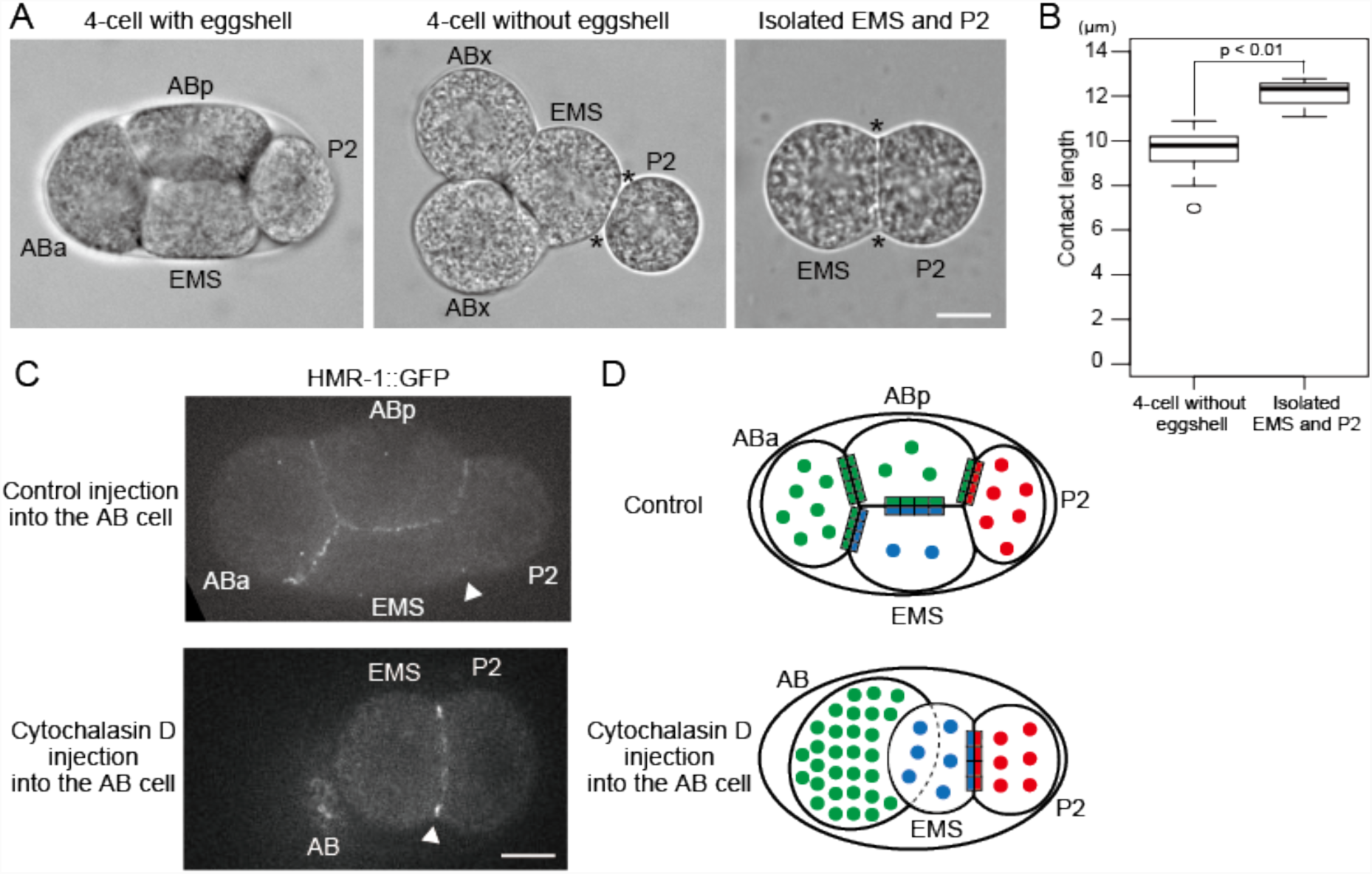
Injection of Cytochalasin D into the AB cell reveals cell-non-autonomous effect on the adhesion strength between EMS and P2 cells. (A) Embryos at the 4-cell stage with and without the eggshell, or without both the eggshell and the AB cell. EMS and P2 cells adhered weakly compared to that with other combinations of cells without the eggshell (middle). Further, removing AB daughter cells strengthened the adhesion between EMS and P2 cells (right). (B) Boxplot of contact length (i.e. the distance between the asterisks in (A)) between EMS and P2 cells with (*n* = 10) or without (*n* = 6) EMS–ABx adhesion. The boxes show the 25th to 75th percentile range. The lines inside the boxes represent the median. Whiskers extend to the most extreme data point within 1.5 interquartile ranges from the box. The open circle is outlier. (C) Embryos expressing HMR-1 (E-cadherin)::GFP (CAL1851 strain). Arrowheads indicate cell–cell contact site between EMS and P2 cells. The signal at the EMS/P2 border was weak in the control embryos, but was strong when Cytochalasin D was injected into the AB cell. All focal planes of Cytochalasin D-injected embryo are shown in Video 4. ‘Control’ means injection of the solution without Cytochalasin D into the AB cell. (D) Schematic diagrams show the localization pattern of cadherins in control and Cytochalasin D-injected embryos. Circles show cadherin molecules within the cytoplasm and squares at the cell borders show cadherin molecules engaged in cell adhesion. Green, blue, and red colors indicate E-cadherin molecules in AB, EMS, and P2 cells, respectively. Note that the total number of circles and squares in each cell is constant. We propose that the EMS cell adheres strongly to the P2 cell in Cytochalasin D-injected blastomeres because the EMS cells does not adhere properly to the AB cell in this condition; thus, surpluses in E-cadherin participate in enhanced EMS–P2 adhesion. Scale bars: 10 μm.

To demonstrate the cell-non-autonomous effect of ABa/p on the strength of adhesion between EMS and P2, we injected Cytochalasin D into the AB cell (the mother of ABa/p cells) to disrupt the cortical integrity of the cells, including the adhesion function. As expected, the injected AB cell did not divide and E-cadherins (cell adhesion molecule) were no longer detected on the surface (Video 4). Consistent with our hypothesis, in this condition, EMS and P2 cells adhered strongly to each other and an increased E-cadherin signal was detected on the border of EMS and P2 cells (Fig. 6C). The result demonstrated that loss of adhesion between EMS and ABa/p cells leads to enhanced adhesion between EMS and P2 cells.

To account for the cell-non-autonomous effect, we propose a limited pool model (Fig. 6D), in which the amount of cell adhesion molecules such as E-cadherin is limited in EMS cells. The majority of the limited pool is normally used for its adhesion to ABa/p cells. As a result, the EMS cell adheres to the P2 cell weakly. Our experiment demonstrated that, for the normal distribution of E-cadherin and weak adhesion at the EMS-P2 border, physical attachment between EMS and ABa/p cells is not sufficient, but an intact AB cell cortex is required. E-cadherin molecules are known to be distributed asymmetrically in 1-cell stage embryos, such that they are enriched in the AB cell but existed at low levels in the P1 cell (the mother of EMS and P2 cells) (Munro *et al.*, 2004; Yamamoto and Kimura, 2017). The limited pool model with the asymmetric distribution of E-cadherin explains the underlying mechanism for the asymmetric attraction between the blastomeres demonstrated in the previous study (Yamamoto and Kimura, 2017).

## Discussion

Previously, direct microinjection into the *C. elegans* embryo was possible only by using carbon-coated quartz needles (Brennan *et al.*, 2013). Unfortunately, this method is restrictive for most researchers due to the special equipment needed to coat the needle. In this study, we made microinjection possible by using uncoated glass needles that are available for most researchers. Direct substance delivery was demonstrated by injecting fluorescent dextran or microbeads in a time- and location-specific manner. When the microinjection was performed at the 1-cell stage, ∼70% of the cells (that fulfilled the leakage criteria) divided at least twice and greater than 40% of the embryos hatched to become larvae. The 1-cell stage embryo seems to be fragile compared to the later stage embryos as (i) the final modifications of the eggshell that protect the embryo are completed after a few mitotic divisions (Stein and Golden, 2015), and (ii) the rate of cell division upon eggshell removal is low at this stage based on our experience. Therefore, the high success rate for the 1-cell stage implies that our method is applicable for later embryonic stages.

The differences between the microinjection method using carbon-coated quartz needles (Brennan *et al.*, 2013) and that using the glass needle in this study are summarized as follows. First is the availability of the needles. The carbon coating of the quartz needles requires special equipment inaccessible for most biology laboratories, whereas the glass needles can be made using an ordinary pipette puller. Second, the arrangement of the embryo, the injection needle, and the holding pipette seems to be more restricted for the glass needle (Fig. 2). Injection with glass needles requires precise alignments between the holding pipette (‘hold-needle alignment’) and the glass needle; moreover, the long axis of the ellipsoidal embryo needs to be perpendicular to the axis of the holding pipette and the glass needle (‘embryo-needle alignment’). In contrast, such strict conditions seemed not to be required for microinjection using the carbon-coated quartz needle, as the carbon-coated quartz needle could be inserted into an embryo that is immobilized at the posterior cortex by the holding pipette (Brennan *et al.*, 2013). The advantage of the carbon-coated quartz needle over the glass needle might not be its hardness to penetrate through the eggshell, but its grip to the surface of the eggshell to avoid slippage. The reason as to why the carbon-coated quartz needle has better grip is unclear.

We could not compare the invasiveness of the two methods. In this study, we quantified the success rates of cell division and hatching. In contrast, there was no such description in the previous report, whereas the authors stated that the injection itself does not inhibit early embryogenesis (Brennan *et al.*, 2013). Considering the reasonable success rate of cell division and hatching with the glass needle, we think our method is sufficiently useful for microinjection in cell and developmental studies. The previous report also found that the carbon-coated quartz needle can be used repeatedly for injection because it is hard. The glass needle can also be used repeatedly for injection, at least three or four times, indicating the glass needle is hard enough for microinjection experiments.

Various experiments involving microinjection approaches have now become possible for *C. elegans* embryos in ordinary biology labs. This approach can also be easily applied to other nematode species with similar eggshells. In this study, we demonstrated that by injecting a fluorescent dye into a blastomere, we could detect the AB cell descendants easily up to adult stages. We also demonstrated a cell-non-autonomous control mechanism of cell adhesion; specifically, inhibiting actin in one cell (AB) at the two-cell stage influenced adhesion between daughter cells of the other cell (P1). Microinjection methods have been used for various experiments using many cell types including mouse oocytes, HeLa cells, and *Xenopus* eggs. Such experiments have now become possible for nematode embryos. For example, microbeads coated with DNA can induce an ectopic polar body-like structure in mouse oocytes (Deng and Li, 2009), induce a nuclear envelope-like structure and avoid autophagy in HeLa cells (Kobayashi *et al.*, 2015), or induce the assembly of microtubules and a bipolar spindle in *Xenopus* egg extracts (Heald *et al.*, 1996). As another example, microbeads coated with Aurora kinase A were reported to act as an artificial centrosome in *Xenopus* egg extracts, and their role in cell division has been investigated (Nguyen *et al.*, 2014). It will be interesting to inject microbeads, in which the surface is functionalized in different ways. We expect that the combination of the microinjection method with sophisticated genetics of *C. elegans* will be a powerful approach to drive cell and developmental biology.

## Materials and Methods

### Strains and maintenance of *C. elegans*

N2 (Bristol), CAL1041 (*oxIs279* [*pie-1p*::GFP::*his-58* + *unc-119*(+)]; *ltIs38*[pAA1; *pie-1p*::GFP::PH(PLC1_delta1_) + *unc-119*(+)]), and CAL1851 (*hmr-1*(cp21[*hmr-1*::GFP + LoxP]) I; *wjIs108* [*pie-1p*::mCherry::*his-58*::*pie-1*_3’UTR + *unc-119*(+)]) strains were used in this study. N2 was used as the wild type. CAL1041 was obtained by mating strains EG4601 and OD58 and CAL1851 was obtained by mating strains LP172 and CAL941 (unc-119 (ed3); *wjIs108* [*pie-1p*::mCherry::*his-58*::*pie-1*_3’UTR + *unc-119*(+)]). These strains were maintained using standard procedures (Brenner, 1974).

### Preparation of glass needles and holding pipettes

Three types of glass needles with different inner diameters (*φ*100, *φ*150, and *φ*660, Fig. 1), and holding pipettes were prepared from a GD-1 glass capillary (Narishige, Tokyo, Japan) using a P-1000 micropipette puller (Sutter Instruments, Novato, CA, USA). The *φ*100-type needle was created by pulling twice with a parameter set of (heat [H], pull [Pu], velocity [V], time [T], pressure [Pr]) = (837, 100, 8, 250, 500) and once with (H, Pu, V, T, Pr) = (837, 100, 15, 250, 500). The *φ*150-type was created by pulling once with (H, Pu, V, Delay, Pr) = (788, 40, 80, 200, 500). The *φ*660-type was created by pulling twice with (H, Pu, V, T, Pr) = (837, 60, 8, 250, 500) and once with (H, Pu, V, T, Pr) = (837, 60, 15, 250, 500). The holding pipette for embryo immobilization was created by pulling once with (H, Pu, V, T, Pr) = (869, 0,150, 200, 200). To create holding pipettes with an inner diameter of 15–25 μm (which is suitable to immobilize the embryo), the taper region was cut by the “glass-on-glass” method (pipette cookbook, Sutter). The tip of the holding pipette was fire-polished using a microforge, MF-900 (Narishige) to create a smooth surface. Curve structures were prepared both in the holding pipette and glass needle at a position 500 μm from the tip using the microforge (Fig. 2, lower images). The curved structures were created by bending the respective capillaries for 15–25 degrees by applying heat from one side of the capillary using the microforge. A brake structure, to avoid acute aspiration, was added to the holding pipette at a region 300 μm from the tip using the microforge. The brake structure was created by leading the holding pipette into a loop of platinum wire and heating it uniformly under the microforge.

### Observation of glass needle tip diameter by SEM

The tips of the glass needles were observed and their tip diameters were measured by SEM (JSM-7500F; JEOL, Tokyo, Japan). Tip segments of the glass needle were carefully cut out using tweezers and were put on the specimen mount for SEM. To measure not only the outer diameter but also the inner diameter, needle tips were tilted on the mount. They were then coated with a 1.5–2-nm thick layer of osmium using a hollow cathode plasma coater (HPC-1S; Vacuum Device, Mito, Japan), to confer a conductive property for observation, and then observed by SEM.

### Measurement of ejection volume (Fig. 1CD, Table 1)

Texas Red-conjugated dextran (1.25 mg/ml, MW = 3000; Invitrogen, D3329; Carlsbad, OR, USA) was loaded into the glass needles using a microloader (Eppendorf, Hamburg, Germany) and dextran was ejected into a glycerol droplet in the same manner as performed in a previous report (Brennan *et al.*, 2013). The injection pressure, injection time, and compensation pressure of a Femtojet (Eppendorf) microinjector were set to 1,000 hPa (14.5 psi), 0.5 s, and 69 pc, respectively. After waiting approximately 1 s until the ejected fluorescent dextran changed to a spherical shape, a generated fluorescent sphere was recorded as a digital image. The diameter of the fluorescent sphere was measured using ImageJ (National Institutes of Health, Bethesda, MD, USA), and the ejection volume was estimated from the diameter. An inverted microscope (Axiovert-100; Zeiss, Oberkochen, Germany) equipped with a 10× PH1-ACHROSTIGMAT 0.25 NA objective (Zeiss) was used to measure the ejection volume. Digital images were obtained using a CCD camera (ORCA C4742-95; Hamamatsu Photonics, Hamamatsu, Japan) controlled by iVision software (BioVision Technologies, Exton, PA, USA).

### Embryo immobilization and microinjection

To manipulate the holding pipette, a coarse micromanipulator (ONM-1; Olympus, Tokyo, Japan) and a fine micromanipulator (ON-2; Olympus) were used. A pneumatic microinjector (IM-11-2; Narishige) was used for embryo immobilization. For the glass needle, the MN-4 coarse manipulator (Narishige) and the MMO-203 fine micromanipulator (Narishige) were used. A pneumatic microinjector (FemtoJet; Eppendorf) was used for injection. These instruments were equipped on an inverted microscope (IX71; Olympus). To align the holding pipette and the glass needle (‘hold-needle alignment’), the tip parts were first visually aligned from a side view (i.e. x-z view, Fig. 2AB). Subsequently, the alignment was examined from a top view (i.e. x-y view, Fig. 2B) visually and under the microscope. After the alignment, the holding pipette and the glass needle were transiently raised toward the z-axis during embryo mounting. The 1-cell stage embryo was cut from an adult and placed in 0.8× EB (1× EB: 118 mM NaCl, 48 mM KCl, 2 mM CaCl_2_, 2 mM MgCl_2_) and transferred to a 24 × 55-mm coverslip (Matsunami, Osaka, Japan); it was then mounted on the inverted microscope. The embryo was immobilized to the holding pipette by applying a negative pressure using a configuration in which the long axis of the embryo was perpendicular to the holding pipette (Fig. 2A, enlarged view at center, Fig. 2B, ‘embryo-needle alignment’). Subsequently, the holding pipette, the glass needle, and the central plane of the embryo were all set on the same focal plane. The objective lens was switched to 100× UPlanSApo 1.40 NA and the glass needle was slowly inserted into the embryo. When injecting a solution into the embryo, injection pressure was applied after the tip of the needle was inserted approximately 2-3 μm across the cell membrane. After the introduction of the substance, the glass needle was first withdrawn by half of the inserted distance, and then the remaining half was withdrawn after approximately 5 s to ensure that the cytoplasm of the embryo did not leak. To test hatching, the injected embryo was transferred to SGM (Shelton and Bowerman, 1996) after it reached the 4-cell stage and was incubated overnight at 25 °C.

### Observation of *C. elegans* embryos, larvae, and adults

Observation of the embryos except Video 2, Fig. 5I (Video 3), and Fig. 6 (Video 4) was performed using a CSU-10 spinning-disk confocal system (Yokogawa, Tokyo, Japan) mounted on the injection microscope (IX71, Olympus) at room temperature. Digital images were obtained using a CCD camera (iXon; Andor Technology, Belfast, UK) controlled by IPLab software (BD Biosciences, Tokyo, Japan). Observation of the embryos in Video 2, Fig. 5I (Video 3), and Fig. 6 (Video 4), larvae and adults, was performed using a CSU-X1 confocal system (Yokogawa) with another IX71 microscope with an iXon CCD camera controlled by MetaMorph imaging software (Molecular Devices, Sunnyvale, CA, USA). For signal observation after hatching, the worms were transferred in a drop of M9 buffer containing 1 mM levamisole (L9756, Sigma, Saint Louis, MO, USA) and were placed on a 26 × 76-mm coverslip (custom-made, Matsunami). Image analyses were conducted using ImageJ software. Images were converted into 8-bit images after brightness and contrast were adjusted using the ‘auto’ setting of the software. To display the entire body of larva or adult stage specimens with a single image, several images were stitched together using the MosaicJ, which is a plugin for ImageJ.

### Fluorescent dyes and microbeads used for substance introduction

Three types of dextran, at different molecular weights (MWs), and four types of microbeads, at different diameters, were examined in this study (Table 1). Texas Red-conjugated dextrans were used as follows: MW = 3,000 (Invitrogen, D3329), MW = 10,000 (Invitrogen, D1863), and MW = 70,000 (Invitrogen, D1863). Microbeads were as follows: 15–20 nm in diameter (Qdot 605, Invitrogen, Q10103MP), 25 nm (micromer®-redF, Micromod 30-00-251; Rostock, Germany), 50 nm (micromer®-redF, Micromod, 30-00-501), and 100 nm (micromer®-redF, Micromod, 30-00-102). Three types of dextran were dissolved in 0.8× EB to a final concentration of 1.25 mg/ml. The Qdot sample was diluted 5-fold from the original solution with water. The concentration of each Micromod microbead solution suspended in water was 10 mg/ml. These solutions including each substance were filled in the tip of the glass needle from the back end with a microloader, and then the glass needle was connected to a Femtojet. If the solution was clogged at the tip of the glass needle, aggregation was resolved using both filtration (25CS020AS, Toyo Roshi, Tokyo, Japan) and sonication (Q125; QSONICA, Newtown, CT, USA). Sonication was performed for 5 s at 70% amplitude with a 10 s rest; these treatments were repeated five times.

### Eggshell removal and blastomere isolation

Eggshells were removed using the method of the previous report (Yamamoto and Kimura, 2017). Embryos were treated with Kao bleach (Kao, Tokyo, Japan) mixed with 10 N KOH at a 3:1 ratio for 90 s, and then transferred to SGM three times to wash the bleaching mixture. The vitelline membrane was removed using a mouth pipette with an approximate 30-40 μm tip inner diameter made from the GD-1 glass capillary (Narishige). To isolate the embryo into two blastomeres at the 2-cell stage, the eggshell-removed embryo was divided into two blastomeres (AB and P1) by manual handling using an eyelash in SGM.

### Statistical comparison of the contact length (Fig. 6B)

From the microscopy images, the length of contact area between EMS and P2 was measured using ImageJ. Mann-Whitney U test was used to compare mean values. For the analyses, R (www.r-project.org) was used. The experiments were not randomized, and the investigators were not blinded to allocation during experiments and outcome assessment.

### Cytochalasin D injection into AB cells of 2-cell stage embryos

20 μg/ml Cytochalasin D (C8273, Sigma) in DMSO and 2.5 mg/ml Texas Red-conjugated dextran (MW = 3000) in water were mixed at a 1:1 ratio to make a 10-μg/ml Cytochalasin D, 1.25-mg/ml dextran solution. This solution was injected into the AB cells of 2-cell stage embryos of CAL1851 strain. ‘Control injection’ in Fig. 6 means injection of a 1:1 mixture of DMSO and 2.5 mg/ml Texas Red-conjugated dextran in water into the AB cell. Dextran was used as an injection marker, and the specific incorporation of Texas Red-conjugated dextran only into the AB cell was confirmed by fluorescence imaging using the spinning-disk confocal system.

## Acknowledgements

*C. elegans* strains CAL1041 and CAL1851 were prepared by Drs. Kenji Kimura and Kazunori Yamamoto (National Institute of Genetics), respectively. We thank Dr. Kazunori Yamamoto for discussion, Dr. Emiko Suzuki (National Institute of Genetics) for teaching SEM operation, Dr. Yukihisa Hamaguchi (Tokyo Institute of Technology) for advice on the micromanipulation system, Drs. Takeshi Itabashi, Shin’ichi Ishiwata (Waseda University), Kenji Ishikawa (Nagoya university), and Shinji Watanabe (Kanazawa University) for technical help and advice on the carbon-coating of quartz needles. We also thank the members of our laboratory for their valuable comments and support. Y.K. is an NIG Postdoctoral Fellow. This work was supported by JSPS KAKENHI grant numbers JP15H04372, JP15KT0083, JP16H00816, 18H05529, and 18H02414.

## Supplementary Videos

**Video 1.** Embryonic cells divide and the embryo hatches after microinjection. The *C. elegans* embryo was injected with egg buffer (EB) at the 1-cell stage. The embryo was the same cell as that shown in Figure 3C. The movie shows a dorsal view of the embryo from a single focal plane. These images were converted into 8-bit images and then converted into the movie after brightness and contrast were adjusted using the ‘auto’ setting of ImageJ software. The top left number indicates the elapsed time after the initial observation (min:sec).

**Video 2.** Location-specific injection into 2-cell stage embryos. The *C. elegans* embryo was injected with egg buffer (EB) and dextran into the AB cell at the 2-cell stage. Note that this embryo is different from the individual shown in Figure 4B. The embryo was transferred from the injection microscope to the CSU-X1 confocal microscope system, and thus the movie begins from the end of 2-cell stage. The movie shows a dorsal view of the embryo. Six different focal planes were stacked at each time point (Z interval = 5 μm). These stacked images were converted into 8-bit images and then converted into a movie after the brightness and contrast were adjusted (as performed for Video 1). Dextran was inherited in the AB cell lineage up to the 8-cell stage. The top left number indicates the elapsed time after starting the observation (min:sec).

**Video 3.** Distribution of fluorescent dextran signal within the embryo. This is the same stage embryo as Figure 5I. Confocal fluorescence images of the dextran-injected embryo were taken every 0.5 μm along the z-axis at the same time point. These images were converted into 8-bit images and then converted into a movie after the brightness and contrast were adjusted (as performed for Video 1). Half of the embryonic cells maintained the fluorescent dextran signal.

**Video 4.** Cytochalasin D injection into the AB cell results in strong adhesion between P1 daughter cells. Confocal fluorescence images of the embryo from Fig. 6C (lower), which was injected with Cytochalasin D into the AB cell at the 2-cell stage. The images were taken every 4 μm along the z-axis at one time point. These images were converted into 8-bit images and then converted into a movie after the brightness and contrast were adjusted (as performed for Video 1). The Cytochalasin D-injected AB cell did not divide and was positioned next to the EMS cell. E-cadherin was no longer at the cortex of the AB cell. EMS and P2 cells strongly adhered to each other as compared to that observed in the control embryo (Fig. 6C, upper).

## Abbreviation

EB: egg buffer
SEM: scanning electron microscopy
SGM: Shelton’s growth medium

